# Targeting CRM1-HMGB1 Nuclear Translocation in Type 2 Diabetes-Driven Metabolic Dysfunction Associated Steatotic Liver Disease

**DOI:** 10.1101/2025.04.09.648024

**Authors:** Prabu Paramasivam, Gabriela Martinez Bravo, Brittany Coffman, Jaya Rajaiya, Satdarshan Paul Monga, Roberto Ivan Mota Alvidrez

## Abstract

Metabolic Dysfunction-Associated Steatotic Liver Disease (MASLD) prevalent in Type-2 Diabetes (T2D) and contributes to progression of Non-Alcoholic Steatohepatitis (NASH). Acetyl-High Mobility Group Box 1 (HMGB1,) a proinflammatory isoform of HMGB1, is released as a DAMP in T2D associated hepatic inflammatory condition. Chromosomal Maintenance 1 (CRM1), a nuclear transporter protein, maintains the nuclear-cytoplasm translocation of hepatic HMGB1 in T2D. We hypothesize that inhibition of CRM1/HMGB1 nuclear shuttling is a therapeutic target for MASLD in T2D. Methods: We performed immunohistochemical analysis of acetyl-HMGB1 and CRM1 in human liver biopsies from control, T2D, and T2D-NASH (n:4 per group). H&E staining to evaluate inflammation and disease stratification. In vitro, studies involved targeted inhibition of CRM1 using Leptomycin-B and HMGB1 with Glycyrrhizin in T2D Huh7 human hepatocytes. Results: T2D subjects with NASH exhibit an increase in acetyl-HMGB1 nuclear and cytoplasmic translocation compared to DM and controls. Acetyl-HMGB1 increased 2-fold in the nucleus and 4-fold in the cytoplasm; CRM1 increased 6-fold in the nucleus and 8-fold in the cytoplasm of T2D/NASH subjects compared to controls. Targeted inhibition of CRM1 and HMGB1 prevented acetyl-HMGB1 hepatocyte release with the most prominent effect in T2D-NASH conditions. Conclusions: Hepatic CRM1/HMGB1 inhibition could be a potential therapeutic target for T2D-driven MASLD.

## INTRODUCTION

Type 2 Diabetes(T2D), a metabolic disorder worldwide(1). It is mainly caused by defective insulin production and pancreatic β-cells and the incapability of insulin-sensitive tissues, such as skeletal muscle, adipose tissue, and liver, to respond to insulin(2). Subsequently, it leads to hyperglycemia, and hepatic tissue crucial role in glucose and lipid homeostasis(3, 4). This results in increased free fatty acid (FFA) release from adipose tissue and hepatic triglyceride synthesis, leading to lipid accumulation in hepatocytes(5). This condition was previously known as hepatic steatosis or Non-Alcoholic Fatty Liver Disease (NAFLD)(6). T2D individuals are at a higher risk of developing NASH due to the close relationship between insulin resistance, lipid metabolism, and inflammation(7). The presence of NASH in T2D subjects significantly increases the risk of liver-related morbidity and mortality(8). The liver inflammation observed in NASH also contributes to systemic inflammation, exacerbating insulin resistance and creating a vicious cycle that worsens both T2D and NASH(9). Reducing liver inflammation is a key therapeutic strategy in managing NASH in T2D subjects(10). This can involve lifestyle changes, pharmacological interventions aimed at reducing insulin resistance or targeting specific inflammatory pathways(11). Addressing insulin resistance in T2D, lipid metabolism, and inflammation simultaneously can help prevent the progression from NAFLD to NASH in T2D subjects and improve overall outcomes(12). MASLD is the latest nomenclature for steatotic liver disease associated with metabolic syndrome directly driven by T2D(13). The increasing prevalence of MASLD worldwide substantially burden on public health(14). MASLD covers a range of liver conditions, from simple steatosis to metabolic dysfunction-associated steatohepatitis (MASH) and cirrhosis(15). Fast paced disease progression of MASLD is a severe concern, so there is an urgent need to identify specific biomarkers and therapeutic targets(15).

While simple steatosis (fat accumulation in the liver) can be relatively non-malicious, it can progress to NASH when combined with inflammation and liver cell damage, a situation that also demands immediate attention(16, 17). In response to the above triggers, immune cells such as Kupffer cells (liver macrophages), recruited monocytes, and other innate immune cells activate in the liver(18). These cells release higher amounts of pro-inflammatory cytokines and chemokines, further amplifying the inflammatory response. The persistent inflammatory environment causes hepatocyte injury and apoptosis(19, 20). Damaged hepatocytes continuously release damage-associated molecular patterns (DAMPs) like HMGB1, further activating immune cells(21). Chronic inflammation leads to the activation of hepatic stellate cells, which produce extracellular matrix proteins, resulting in liver fibrosis-a hallmark of NASH(22, 23). In our previous studies, we have shown that HMGB1 alleviates hyperglycemia severity and enhance glucose tolerance. We revealed a substantial reduction in glucose and enhanced glucose tolerance in HMGB1 knockout diabetic mice. In addition, RNA sequence data showed differential genes and molecular pathways in liver and skeletal muscle, suggesting the potential of HMGB1 knockdown in inducing phenotypic changes in hyperglycemic mice(24). Importantly, HMGB1 silencing appears to have key roles in protecting from liver inflammation. HMGB1 exhibits high homology across species, with 100% similarity between mice and rats and 99% between rodents and humans(25). In humans, HMGB1 is composed of 215 amino acids organized into two DNA-binding domains, the HMG A box (amino acids 9–79) and the HMG B box (95–163), followed by a C-terminal acidic tail (186–215) and a short functional N-terminal region(26-29). HMGB1’s stable nuclear localization is maintained by two nuclear localization signals, NLS1 (amino acids 28–44) and NLS2 (179–185), while its nuclear export signal (NES) is embedded within the DNA-binding domains(30). Under oxidative stress, post translational modifications (PTMs) like acetylation and deacetylation, mediated by histone acetyltransferases and deacetylases, enable HMGB1 to shuttle between the nucleus and cytoplasm by preventing nuclear re-entry through the NLS(31). Once acetylated, HMGB1 binds to CRM1 (also known as Exportin 1), which exports it to the cytoplasm(32-34).

Liver inflammation in T2D drives disease progression towards MASLD, exacerbating systemic insulin resistance, creating a cycle that can lead to severe liver and metabolic complications(35-37). Because of the role of HMGB1 in glycemic control, insulin resistance and chronic inflammation in T2D, we propose that targeting HMGB1 may offer a therapeutic strategy for restoring glucose balance in T2D-driven MASLD. The main aim of our study is to examine the translocation of CRM1 and acetyl-HMGB1 in nuclear/cytoplasmic translocation in human liver tissues from T2D and T2D/NASH, compared to healthy livers. Our experimental hypothesis is that silencing CRM1 will limit acetyl-HMGB1 release from the nucleus to the cytoplasm in our T2D-insulin resistance Huh7 in vitro model to validate CRM1/HMGB1 nuclear export as a therapeutic target for T2D-driven MASLD.

## MATERIALS AND METHODS

### Human liver samples

Formalin-fixed paraffin-embedded human liver biopsies from control, T2D, and T2D/NASH human subject were provided by Dr. Paul Monga at the Pittsburgh Liver Research Center from the University of Pittsburgh in agreement with a shared collaboration agreement with the University of New Mexico. All procedures were approved by the IRB at the University of Pittsburgh in agreement with the University of New Mexico.

### Immunohistochemistry analysis of acetyl-HMGB1 and CRM1

Liver sections of formalin-fixed paraffin-embedded tissue (FFPE) are cut at 4-5 microns, mounted on charged (+) slides, and baked at 60°C for 60 minutes. The Ventana Discovery platform was used for deparaffinization staining. Prior to antibody application, slides were treated with Discovery CC1 (Ventana 950-500) for 24 minutes at 100°C. CRM1 antibody (Abcam ab24189) was diluted to 1:200, and HMGB1 [Acetyl-Lys12] antibody (Aviva Sys Bio OASG03545) was diluted to 1:100 in Discovery P.S.S. Diluent (Ventana 760-212). Both antibodies were hand applied and incubated for 60 minutes at 36°C. Following antibody incubation, the slides were performed with secondary antibody anti-Rb HQ (Ventana 760-4815) for 12 minutes and conjugate anti-HQ HRP (Ventana 760-4820) for 12 minutes. Detection was achieved using Ventana DAB CM chromogen (Ventana 760-4304). Slides are counterstained with Hematoxylin (Ventana 760-2021) and Bluing solution (Ventana 760-2037), removed from the auto-stainer, and hand coverslipped. All slides were scanned on the Leica Aperio AT2 digital scanner at 20X magnification. Quantitative analysis of CRM1 and HMGB1 was performed using the HALO analysis and annotation software package (Indica Labs, Albuquerque, NM). Hematoxylin was used as a nuclear counterstain for cell identification while DAB (3,3’-Diaminobenzidine) chromogen staining to identify the antibodies. The HALO system’s Figure Maker feature was used to generate the magnification series of images with and without HALO markup. In the (supplement figure 2 A-H), illustrated, to distinction between cytoplasmic and nucleus region using hematoxylin as a nuclear counter stain in hepatic and vascular region and figure showed masked and unmasked localization of acetyl-HMGB1 and CRM1 using The HALO system’s Figure Maker feature.

### In vitro studies

We purchased a hepatocellular carcinoma Huh7 cell line from ATCC (Manassas, VA, USA). Huh7 cells were maintained at 37 °C and 5% CO2 in DMEM F12 (Gibco, Cat. No. 12634-010) and supplemented with 10% of fetal bovine serum (Gibco, Cat. No. 16000044), 1% L-glutamine (Gibco, Cat. No. 25030-083), and 1% antibiotic/antimycotic mixture (Sigma, Cat. No. A5955). The 10% DMEM media changed every 48 hours. For our different in vitro treatments, confluent Huh7 cells were subject to 10% FBS DMEM with normal glucose (17.5mM) and normal insulin for our non-T2D conditions and for T2D conditions we stimulated cells with high glucose (40 mM) and high insulin (100nM) for 72hr plus the addition of palmitate (0.5 mM) treated for 24h to induce insulin resistance in our T2D in vitro model(38, 39). CRM1 was inhibited by Leptomycin B (LMB, 25nM) for 12hrs, which is a known pharmacological inhibitor for CRM1 used to inhibit release of acetyl-HMGB1 from nucleus to cytoplasm. HMGB1 was inhibited by Glycyrrhizin (GA, 50µM) for 24hrs, which is a known pharmacological inhibitor for HMGB1 used to inhibit acetyl-HMGB1 release.

### Measurements of cell viability (MTT assay)

An MTT assay, a meticulous method, was employed to determine cell viability. Huh7 cells were subjected to palmitate/sodium oleate at various ratios in 96-well plates for 24 h. The cells were then incubated with MTT at a precise concentration for a specific duration at a controlled temperature. The absorbance was measured using a reliable microplate reader. The data, expressed as the mean ± standard deviation, were the result of experiments performed in five replicates, ensuring the robustness of our findings(40, 41).

### Nuclear and cytoplasmic protein extraction

After the treatment, Huh7 cells were harvested, and for nuclear and cytoplasmic extraction, they were prepared using a NE-PER Nuclear Cytoplasmic Extraction Reagent kit (Pierce, Rockford, IL, USA) per the manufacturer’s instructions. Briefly, the treated cells were washed twice with cold PBS and centrifuged at 500g for 3 min. The cell pellet was suspended by vertexing in 200μl of cytoplasmic extraction reagent I. The suspension was incubated on ice for 10 min, followed by the addition of 11μl of a second cytoplasmic extraction reagent II, vortexed for 5 s, incubated on ice for 1 min, and centrifuged for 5min at 16,000 g. The supernatant fraction (cytoplasmic extract) was transferred to a pre-chilled tube. The insoluble pellet fraction containing crude nuclei was resuspended in 100μl of nuclear extraction reagent by vertexing for 15 s and incubated on ice for 10min, then centrifuged for 10min at 16,000 g. Nuclear and cytoplasmic lysates were stored at -80 degrees until needed.

### Immunoblotting for acetyl-HMGB1 and CRM1

The concentrations of total cell lysates and nuclear or cytoplasmic proteins were estimated by the bicinchoninic acid assay (BCA) (Pierce BCA protein assay kit, #23227, Thermo Fisher Scientific, USA). In brief, 20ug of proteins were separated on a 4-12% SDS–polyacrylamide gel. Following electrophoresis, the proteins were transferred to a PVDF membrane. The blotted membranes were blocked for 1h in 5% nonfat dry milk in Tris-buffered saline containing 0.1% Tween 20 and then incubated with the indicated primary antibodies such as acetyl-HMGB1(1:1000; Cat # OASG03545 Aviva Systems biology, San Diego, CA, USA), CRM1(1:1000; Cat # 24189; Abcam) and anti-β-actin antibody (1:1000, Cat #A2228, Sigma-Aldrich), anti-α tubulin antibody (1:1000, Cat #T5168, Sigma-Aldrich) overnight at 4-degree temperature. The blots were washed four times with TBST and incubated for 1 h with horseradish peroxidase-conjugated anti-rabbit IgG antibody (1:12000, Cat #A27036, ThermoFisher Scientific, USA) or anti-mouse IgG antibody (1:2000, Cat #A28177, Thermo Fisher Scientific, USA) in 5% nonfat milk. Protein expression was detected using enhanced chemiluminescence (ELC, #34577, Thermo Fisher Scientific, USA). Blots were stripped and re-probed for loading control with β-actin (cytoplasm) or alpha-tubulin (nuclear protein) to ensure equal loading of protein.

### Total RNA extraction and quantitative Real-Time PCR

Total RNA extracted from Huh7 transfected cells using PureLinK RNA mini Kit (Cat. # 2775172, Invitrogen/Thermo Fisher Scientific, USA). RNA quantity and quality were assessed by NanoDrop 2000 (Thermo Fisher Scientific, USA) instrument. 2 µg of total RNA from each sample was used to convert cDNA using the High-Capacity cDNA Reverse transcription kit (Cat. #. 4368814, Applied Biosystems/Thermo Fisher Scientific, USA) according to the manufacturer’s protocol. Real-time PCR was performed using the 100ng cDNA product, Power SYBR Green PCR Master Mix (Cat. #. 4367659, Applied Biosystems/Thermo Fisher Scientific, USA) and qPCR. Amplification conditions included 2 min at 50°C and 10 min at 95°C, and then, they were run for 40 cycles at 95oC for 15 sec and 60°C for 1 min on the LightCycler 480 II (Roche). HMGB1 and CRM1 gene expression was normalized to housekeeping gene β-actin, and relative gene expression was calculated using 2^-ΔΔCT^ method(42).

### Statistical analysis

All statistical analysis was performed using GraphPad Prism 9. For the immunohistochemistry of acetyl-HMGB1 and CRM1 analysis, human liver tissue sample sizes were an n of 4 per group, ensuring comprehensive coverage. In the HCM experiment, the sample size was n: 4-5 per group/condition, providing a solid basis for our conclusions. Band intensity in immunoblots, n:3 per group per assay. For molecular biology experiments, assays were performed at least 3 times. Data: means ± SEM. We employed one- and two-way ANOVA with Tukey’s post hoc modified t-tests or two-tailed t-test where appropriate. Our findings reached statistical significance at p< 0.05.

## RESULTS

### T2D/NASH livers evidenced increase in hepatic inflammatory infiltrates with advanced fatty-liver disease stage

We examined granulocytes and agranulocyte cell infiltration in liver sections using hematoxylin and eosin staining from control, T2D, and T2D/NASH subjects. Our unbiased morphologic and histologic examination was performed by a licensed and certified clinical pathologist (Dr. Brittany Coffman) who performed unbiased analysis assessing the morphology and degree of steatosis liver tissues in the study groups. The assessment confirmed the control groupA(A-H) has no steatosis, T2DA(A-H)shows infiltrated inflammatory cells between grade 2 and stage 1 steatosis, and T2D/NASH A(A-H) group histological sections appear all between stage 4 steatosis (Fig.1A). We further assessed to distinguish granulocytes and agranulocytes from infiltrated cells using nuclear size. Granulocytes (Basophils, Eosinophils, Neutrophils) are shown as being below 4.4 µM nuclei size and agranulocytes (lymphocytes and monocytes) being above 4.4µM of nuclei size (Fig.1A). To distinguish between hepatic and vascular regions of interest, we analyzed inflammatory immune cell infiltration in both hepatic (Fig. 1A) and vascular regions (Supp. Fig. 1A). Our analysis of total cells per tissue area showed no difference between agranulocytes and granulocytes in both hepatic (Fig. 1B) and vascular regions (Supp. Fig. 1B). Further analysis of immune cell infiltration, specifically the average cytoplasmic area per total cell, revealed a significant increase in granulocytes compared to agranulocytes in both hepatic (Fig. 1C) and vascular regions (Supp. Fig. 1C). Additionally, a significant difference in the cytoplasmic-to-nucleus area ratio was observed in agranulocytes within the hepatic regions of T2D and T2D/NASH liver sections compared to controls (Fig. 1D). In contrast, in the vascular regions, the cytoplasmic-to-nucleus area ratio indicated a significant increase in granulocyte infiltration compared to agranulocytes in controls (Supp. Fig. 1D). These findings suggest that granulocyte infiltration in vascular tissue is minimal and largely restricted to vascular regions. However, in hepatic regions, marked granulocyte infiltration was observed, particularly in cases with moderate to severe steatosis and tissue damage in T2D and T2D/NASH liver sections. This hepatic infiltration was associated with increased inflammation, lipid accumulation, and subsequent liver injury and tissue damage, highlighting a differential inflammatory response between hepatic and vascular tissues.

**Figure 1:**
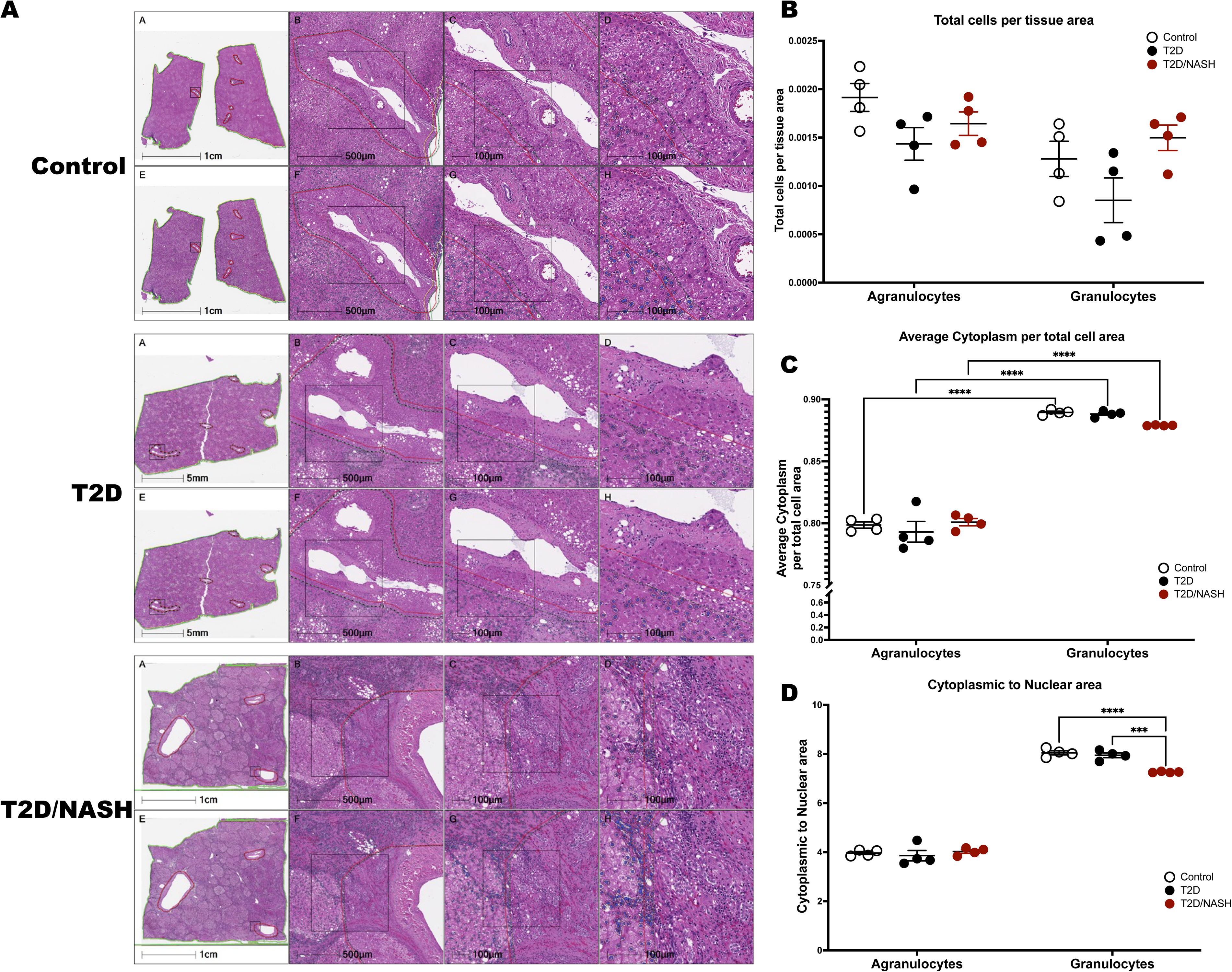
Human liver tissue histological analysis using hematoxylin and eosin staining from control, T2D and T2D/NASH liver samples. We performed analysis of liver section in areas focused on hepatic parenchyma regions of interest without vascular rich areas. A) Control group(A-H): normal histological morphology and no steatosis; T2D(A-H) group shows infiltration of blood cells and steatosis grade 2 and stage 1; T2D/NASH(A-H) group; histological section appears stage 4 and grade 1 steatosis (B-D). Agranulocytes and granulocytes quantification was analyzed and expressed as follows: B) total cell per tissue area, C) average cytoplasm per cell area, and D) average nucleus per total cell area. All data represented mean+/-SEM. All granulocytes and Agranulocytes numbers in control compared to T2D or T2D/NASH of humans (n=4) are used for One-way ANOVA analysis. Significance levels: *:<0.05, **: 0.001-0.01, ***: 0.0001, ****: <0.0001.

### T2D/NASH livers evidence hepatic acetyl-HMGB1 translocation from nucleus to cytoplasm

Comprehensive liver histology assessment showed increasing granulocyte infiltration characteristic of steatosis and inflammation. We next investigated the pro-inflammatory form of acetyl-HMGB1 using immunohistochemistry in the hepatic region of liver tissue of the study groups. We have distinguished acetyl-HMGB1 protein distribution in the nuclei and cytoplasm of liver sections using hematoxylin nuclear counter staining and analysis with HALO. Figure. 2 shows the immunohistochemistry of the acetyl-HMGB1 protein in the liver sections of control (Fig.2A(A-H)), T2D (Fig.2B(A-H)), and T2D/NASH (Fig.2C(A-H)) groups. Quantitation of acetyl-HMGB1 was analyzed based on weak (Fig. 2D), moderate (Fig.2E), and strong positive cells (Fig 2F). Notably, acetyl-HMGB1was particularly increased in cytoplasmic and nucleus region of T2D/NASH group compared to control group (Fig.2F).Additionally, nucleus/cytoplasmic ratio of acetyl-HMGB1 in T2D and T2D/NASH groups showed to be increased compared to control subjects (Fig.2F). In Supp. Fig. 3A-F, we show the vascular region of acetyl-HMGB1 protein nucleocytoplasmic localizations. There were minimal positive stained cells observed even though the hepatic region of hepatocytes cells released acetyl-HMGB1 as a crucial role in inflammation and tissue damage. These finding suggests the pro-inflammatory isoform of acetyl-HMGB1 is more pronounced in cytoplasmic hepatocytes in T2D/NASH, exacerbating liver inflammation and tissue damage.

**Figure 2.**
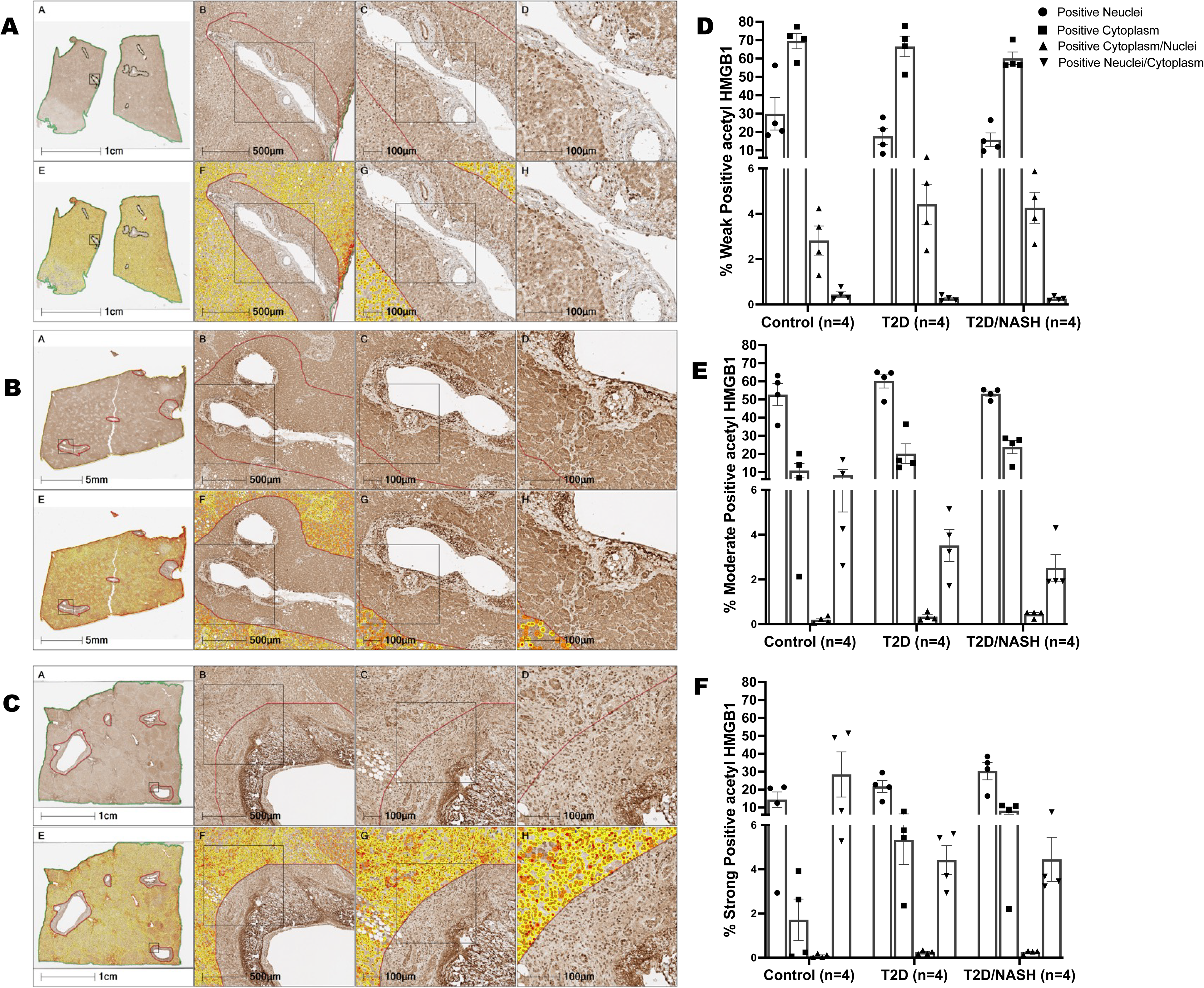
Hepatic acetyl-HMGB1 protein quantification in nucleus and cytoplasmic of the liver tissue from control, T2D, and T2D/NASH human samples. Formalin-fixed paraffin-embedded human liver biopsies from control, T2D, and T2D/NASH human patient samples were used for the analysis of acetyl-HMGB1 protein, which is one of the proinflammatory forms of HMGB1 to distinguish disease severity. Weak positive, moderate positive, and strong positive of acetyl HMGB1 protein was analyzed as follows: A(A-H) Control, B(A-H) T2D, C(A-H) T2D/NASH, D) % weak positive acetyl-HMGB1 E) % moderate positive acetyl-HMGB1 F) % strong positive acetyl-HMGB1 quantification of nucleus, cytoplasm, cytoplasmic/nuclear and nuclear/cytoplasmic ratios of acetyl-HMGB1. All data represented mean+/-SEM. All nuclear and cytoplasmic acetyl-HMGB1 protein in controls compared to T2D or T2D/NASH of humans (n=4) are used for One-way ANOVA analysis. Significance levels: *:<0.05, **: 0.001-0.01, ***: 0.0001, ****: <0.0001. Immunohistochemistry of acetyl-HMGB1 protein color shows as brown and captured at x400.

### Increased nuclear localization of CRM1 protein present In T2D and T2D/NASH liver sections

To assess the nuclear export receptor CRM1, immunohistochemistry was performed in both hepatic and vascular regions. Due to CRM1’s sensitivity, staining posed challenges in our liver sections. CRM1-positive cells were further analyzed based on the percentage of cells with weak, moderate, and strong staining. In both the hepatic (Fig. 3A(A-H), B(A_H)and C(A-H)) and vascular (Supp. Fig. 4A-C) regions, weak CRM1 staining was more prominent than moderate or strong staining. We observed increased weak CRM1 staining in both hepatic (Fig. 3D) and vascular (Supp. Fig. 4D) regions in T2D and T2D/NASH samples compared to controls. However, cells with moderate and strong nuclear CRM1 staining were less than 1% in both hepatic (Fig. 3E-F) and vascular (Supp. Fig. 4E-F) regions across groups. Notably, cytoplasmic CRM1 staining was absent in the hepatic region (Fig. 3), while it was present in the vascular region (Supp. Fig. 4D). This finding highlights the presence of CRM1 staining in hepatocyte nuclei within the hepatic region of T2D/NASH liver sections, representing a novel observation in our study.

**Figure 3.**
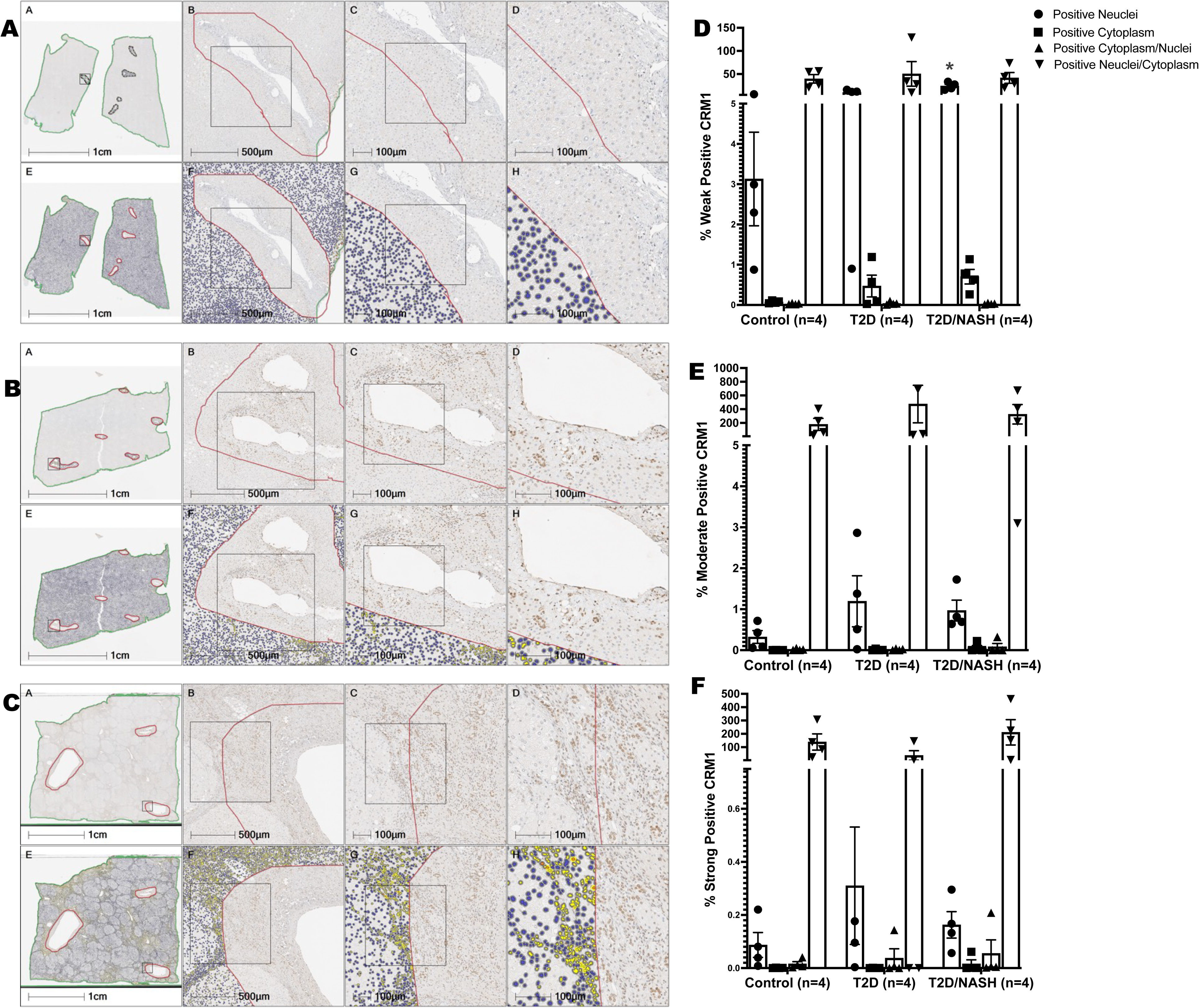
Hepatic CRM1 protein quantification in nucleus and cytoplasmic of the liver tissue from control, T2D, and T2D/NASH human samples. Formalin-fixed paraffin-embedded human liver biopsies from control, T2D, and T2D/NASH human patient samples were used for the analysis of CRM1 protein, which is one of the nuclear exporter proteins. Weak positive, moderate positive, and strong positive of CRM1 protein analyzed as follows: A(A-H) Control, B(A-H) T2D, C(A-H) T2D/NASH, D) % weak positive CRM1, E) % moderate positive CRM1, F) % strong positive CRM1, quantification of nucleus, cytoplasm, cytoplasmic/nuclear and nuclear/cytoplasmic ratios of CRM1. All data represented mean+/- SEM. All nucleus and cytoplasmic acetyl-HMGB1 protein in control compared to T2D or T2D/NASH of humans (n=4) are used for One-way ANOVA analysis. Significance levels: *:<0.05, **: 0.001-0.01, ***: 0.0001, ****: <0.0001. Immunohistochemistry of acetyl-HMGB1 protein color shows as brown and captured at x400.

### In vitro inhibition of CRM1 in fatty acid-induced T2D human hepatocytes limits acetyl-HMGB1 export to cytoplasm

Acetyl-HMGB1 is translocated from the nucleus to the cytoplasm, leading to liver damage, steatosis, and inflammation. We investigated whether inhibiting nuclear transporter CRM1 to limiting acetyl-HMGB1 translocation to cytoplasm. Huh7 cells were treated with leptomycin B, a known CRM1 pharmacological inhibitor, in the presence and absence of T2D and free fatty-induced NASH conditions. Figure 4. Illustrates immunofluorescence analysis of cytoplasmic and nuclear acetyl-HMGB1 and CRM1 (Fig.4C). In Huh7 cells, T2D or T2D/NASH condition induced cytoplasmic acetyl-HMGB1, significantly decreased in T2D/NASH with LMB condition compared to control cells with LMB (Fig.4D-E). On other hand, both cytoplasmic and nuclear CRM1level were decreased in T2D/NASH with LMB condition compared to control cells with LMB (Fig.4 F-G).

**Figure 4.**
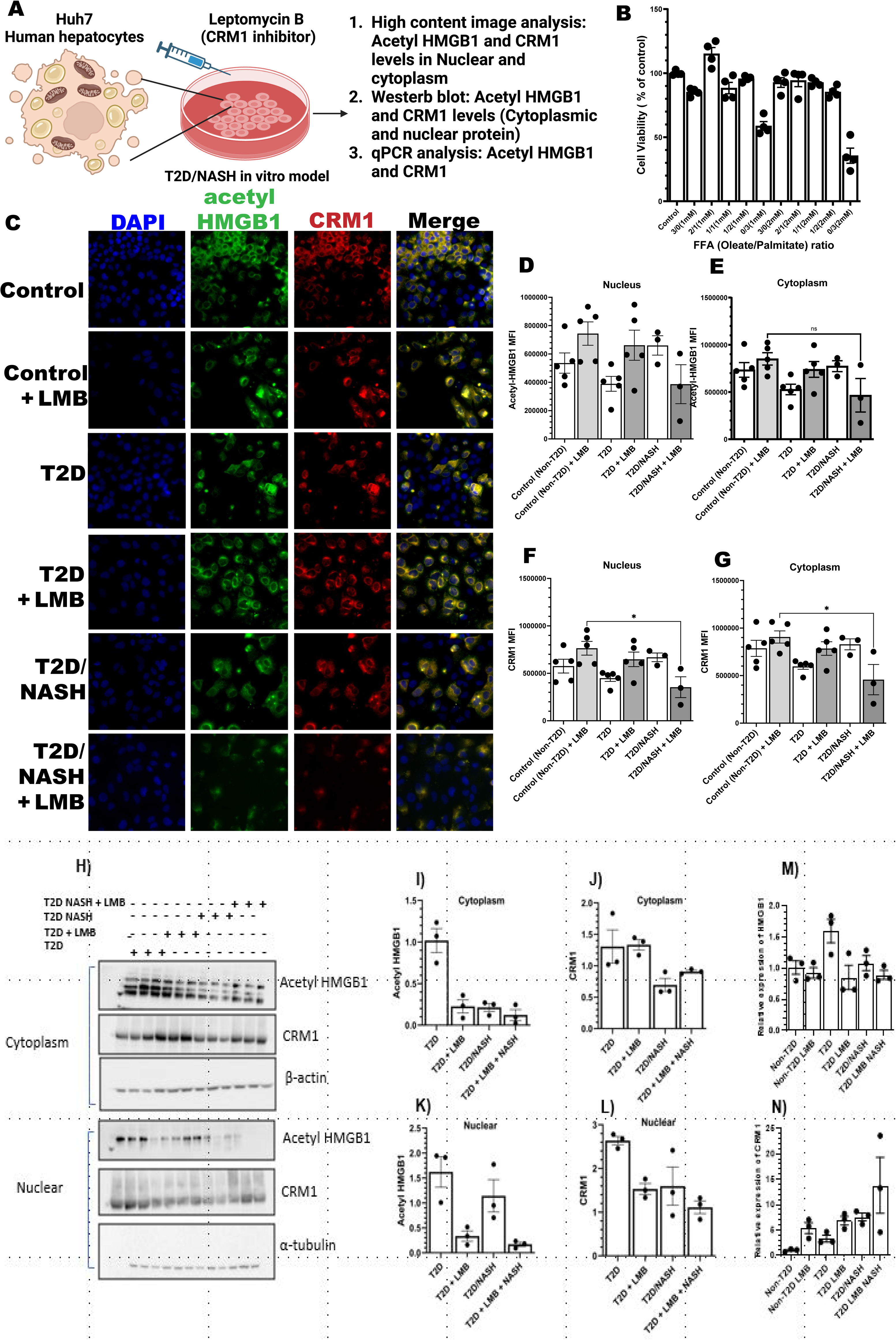
T2D/NASH in vitro model evidenced acetyl-HMGB1 nuclear/cytoplasmic transport in Huh human hepatocytes treated with CRM1 inhibitor. To mimic the T2D/NASH invitro model, Huh7 human hepatocyte cells were treated with high glucose, high insulin, palmitate, and free fatty acid (Oleate/palmitate ratio)-induced T2D/NASH to study acetyl-HMGB1 in the nucleus and cytoplasm. CRM1 was inhibited with LMB in vitro following our methods outlined in Figure 4A. Cell viability MTT assay free fatty acid (sodium oleate/palmitate) different ratio used measure cell viability assay(B). LMB inhibited acetyl-HMGB1 and CRM1 translocation in T2D, T2D + LMB, T2D/NASH and T2D/NASH + LMB conditions (C). Nuclear and cytoplasmic acetyl-HMGB1 and CRM1 imaging quantification (D-G). Western blot analysis of nuclear and cytoplasmic acetyl-HMGB1 and CRM1 protein (H-L). Quantitative Real time PCR analysis of HMGB1 and CRM1 gene expression (M-N). All data represented mean+/-SEM. All nucleus and cytoplasmic acetyl-HMGB1 protein in control compared to T2D or T2D/NASH of humans (n=4) are used for One-way ANOVA analysis. Significance levels: *:<0.05, **: 0.001-0.01, ***: 0.0001, ****: <0.0001.

Additionally, cytoplasmic, and nuclear acetyl-HMGB1 and CRM1 protein levels were further analyzed by immunoblotting. Immunoblotting analysis showed Huh7 cells, cytoplasmic and nuclear acetyl-HMGB1 and CRM1 protein were elevated in T2D or T2D/NASH conditions but were rescued in Huh7 cells treated with LMB in both T2D and T2D/NASH condition (Fig.4I, K). CRM1 protein was retained in nucleus and cytoplasm under LMB treatment with both T2D T2D/NASH conditions, compared to T2D or T2D/NASH (Fig.4J, L). HMGB1 gene expression increased in T2D and T2D/NASH conditions but decreased with LMB treated both T2D and T2D/NASH conditions (Fig.4M). CRM1 gene expression showed increasing trend in LMB treated both T2D and T2D/NASH condition compared to LMB treated control cells (Fig.5N). Nucleocytoplasmic localization of acetyl-HMGB1 and CRM1 in control cells treated with or without Leptomycin B (LMB, 25 nM) show a potential relationship as it seems to be increased in cytoplasmic and nuclear CRM1 along with increased cytoplasmic localization and deceased nuclear HMGB positive staining in treated cells compared untreated control cells (Supp. Fig.5A-E).

**Figure 5.**
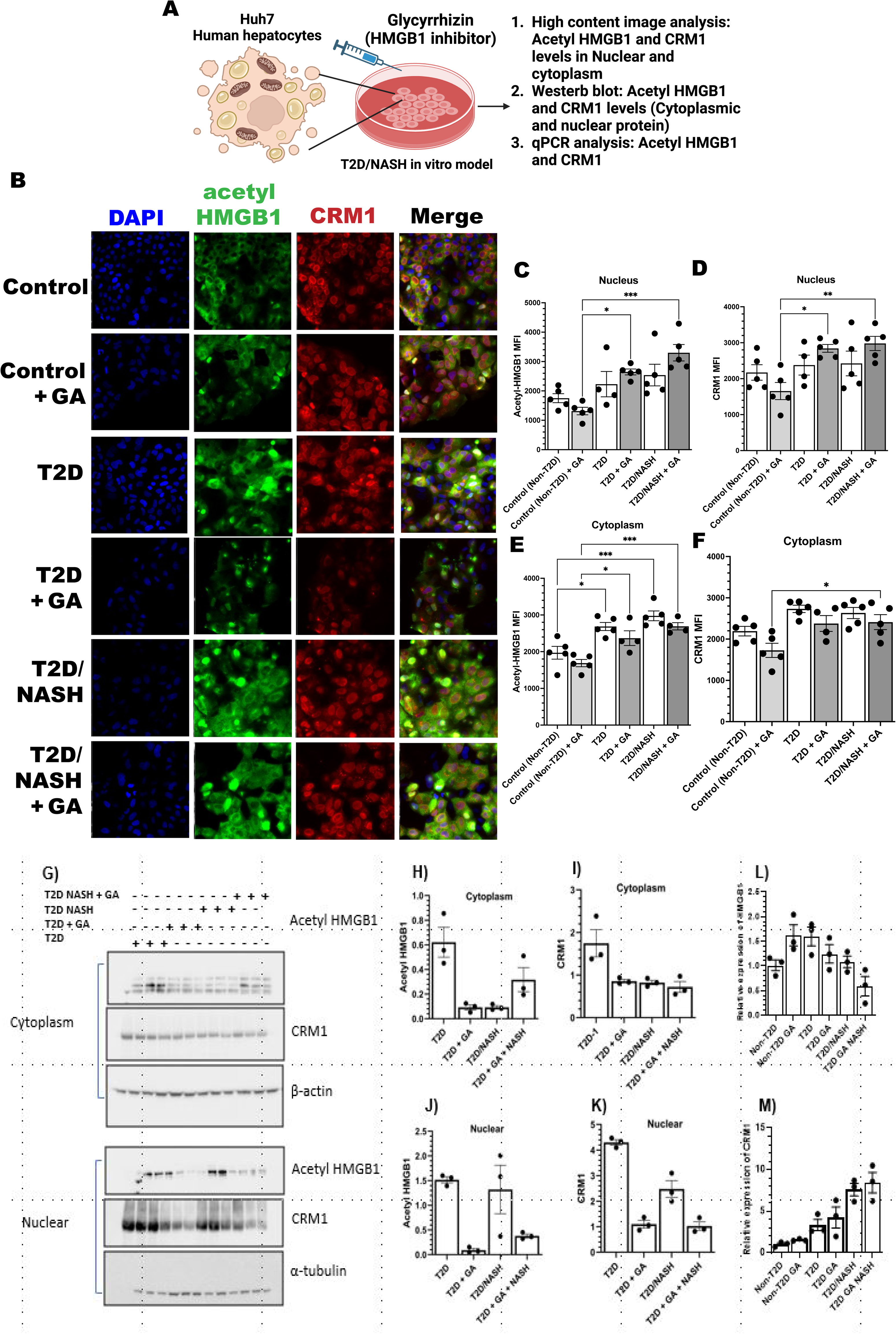
T2D/NASH in vitro model evidenced acetyl-HMGB1 nuclear/cytoplasmic transport in Huh human hepatocytes treated with HMGB1 inhibitor. To mimic the T2D/NASH invitro model, Huh7 human hepatocyte cells were treated with high glucose, high insulin, palmitate, and free fatty acid (Oleate/palmitate ratio)-induced T2D/NASH to study acetyl-HMGB1 in the nucleus and cytoplasm(A). HMGB1 was inhibited with GA. Results show GA treatment and analysis of acetyl-HMGB1 and CRM1 translocation in T2D, T2D+GA, T2D/NASH and T2D/NASH+GA conditions (B-F) with nuclear and cytoplasmic acetyl-HMGB1 and CRM1 imaging quantification. Western blot analysis of nuclear and cytoplasmic acetyl-HMGB1 and CRM1 protein (G-K). Quantitative Real time PCR analysis of HMGB1 and CRM1 gene expression (L-M). All data represented mean+/-SEM. All nucleus and cytoplasmic acetyl-HMGB1 protein in control compared to T2D or T2D/NASH of humans (n=4) are used for One-way ANOVA analysis. Significance levels: *:<0.05, **: 0.001-0.01, ***: 0.0001, ****: <0.0001.

### Inhibition of HMGB1 in fatty acid-induced T2D conditions distinctly mediate acetyl-HMGB1 nuclear translocation

Our previous finding showed how inhibition of CRM1 limits acetyl-HMGB1 release, thereby reducing inflammation and liver damage. Now we moved on to whether acetyl-HMGB1 can be inhibited by the pharmacological inhibitor Glycyrrhizin and collected mRNA, protein, and immunofluorescence of acetyl-HMGB1 and CRM1 un Huh7 cells. Figure 5. illustrates immunofluorescence analysis of cytoplasmic and nuclear acetyl-HMGB1 and CRM1 (Fig.5B). In Huh7 cells, cytoplasmic acetyl-HMGB1 was significantly increased under T2D, or T2D/NASH condition compared to control cells, with further increase observed in GA treated with both T2D and T2D/NASH condition compared to GA treated control cells (Fig.5E).

Similar trend was observed for nuclear acetyl-HMGB1(Fig.5C). However, nuclear CRM1 protein remained unchanged across T2D, T2D/NASH and control condition. Notably, nuclear CRM1 significantly increased in GA treated with both T2D and T2D/NASH condition compared to GA treated control cells (Fig.5D). Cytoplasmic CRM1 levels also significantly increased in GA treated T2D/NASH condition compared to GA treated to control cells (Fig. 5F). Immunoblotting analysis results showed, Huh7 cells, T2D and T2D/NASH induced acetyl-HMGB1 was reduced following GA treatment in both condition (Fig.5J). Nuclear CRM1 level were elevated in GA treated T2D cells compared to T2D condition, but no significant changes observed in T2D/NASH treated cells (Fig.5K). Cytoplasmic acetyl-HMGB1 and CRM1 level inconsistent between T2D and T2D/NASH treated cells (Fig.5I, K). HMGB1 gene expression was increased in T2D and T2D/NASH conditions, but decreased following GA treated both T2D and T2D/NASH conditions but not in GA treated control cells (Fig.5L). CRM1 gene expression showed increasing trend in GA treated both T2D and t2D/NASH condition compared to GA treated control cells (Fig.5M). Supp. Fig.6A-E illustrates the nucleocytoplasmic localization of acetyl-HMGB1 and CRM1 in control cells treated with or without Glycyrrhizin (GA, 50µM) showing a similar pattern with a decrease in cytoplasmic and nuclear acetyl-HMGB1 along with increased cytoplasmic and nuclear CRM1 positive staining in treated cells compared untreated control cells.

## Discussion

Our study has been able to evidence how the CRM1/acetyl-HMGB1 axis is an ideal therapeutic target in T2D-driven MASLD. The strength of our study relies in our data evidencing how granulocyte infiltration, nuclear-cytoplasmic localization of proinflammatory acetyl-HMGB1, and CRM1 in human hepatocytes are prominent in the liver section of T2D/NASH subjects. Additionally, unbiased morphologic and histologic examination conducted by an unbiased certified clinical pathologist, confirmed the presence and grading of steatosis in the liver section from T2D/NASH and T2D compared to control subjects. In our study, MASLD was not definitively diagnosed in all patients, since instead we would be relying on clinical criteria and biomarker analysis for such diagnosis. This estimation approach may not fully capture MASLD’s complexity but allowed us to provide evidence on how our approach can improve targeting based on our results from liver biopsies. Therefore, our findings underline the crucial role of inflammation and fat accumulation, the focus of our study, in the progression of NASH that can evolve to MASLD.

We start our discussion focused on our results showing an increased infiltration of granulocytes compared to agranulocytes in the cytoplasm of hepatic cells. These findings align with a recent study reported that granulocytes, such as neutrophils, infiltrate hepatic liver tissue, a crucial histopathological indication of chronic liver conditions like NASH(43, 44). The abundance of granulocytes like neutrophils are abundant during different stages of MASLD, and various rodent studies suggested a disease-promoting role, especially at the onset of NASH(45). The infiltration of granulocytes has a causal role in human NASH subjects, further highlighting our findings. Proinflammatory HMGB1 is secreted under stress and inflammatory conditions particularly after NF-KB activation. Secreted HMGB1 is often acetylated and acetylated-HMGB1 induce further sterile inflammation via RAGE activation responses through different mechanisms(46-49). In our study, we found increased acetyl-HMGB1 in cytoplasm in T2D/NASH and T2D compared to healthy controls. Recent study revealed the importance of nucleus to cytoplasm translocation of acetyl-HMGB1 in invitro and alcohol-induced liver mouse model(50).Similarly, an animal model study showed reduced systemic HMGB1 levels in high fat-diet fed, HC-HMGB1 knockout mice(51) and Chen et al. 2013 and Gaskell et al. 2018 reported that HMGB1 plays a crucial role in liver damage and chronic liver disease(52, 53) and increased circulating HMGB1 in NAFLD mouse model neutralizing antibodies to HMGB1 was protective(54). Also, oxygen-glucose deprivation/reperfusion-induced hepatic cells showed nuclear-cytoplasmic translocation of acetyl-HMGB1, leading to increased pro-inflammatory cytokines, oxidative stress, and apoptosis(55). These findings indicate hepatic HMGB1 signaling major role in liver damage and chronic liver diseases.

Based on our findings in human liver sections we decided to perform targeted inhibition in vitro in human hepatocytes using Glycyrrhizin (GA), an extract of licorice root, a triterpenoid saponin glycoside with a potential role in immunomodulating, anti-inflammatory, hepato-and neuro-protective, and antineoplastic activities(56-58). Previous studies have shown that Glycyrrhizin suppresses HMGB1 expression and mitogenic activity, impeding its DNA-binding function(56). Sakamoto et al. initially employed biochemical techniques to show that Glycyrrhizin binds directly to HMGB1, inducing conformational changes that prevent its DNA-binding activity(59). Later, Mollica et al. confirmed this binding using nuclear magnetic resonance and fluorescence techniques, demonstrating that Glycyrrhizin docks into the DNA-binding concaves of both HMGB1 boxes (56). These studies confirm that the protective effects of Glycyrrhizin are associated with the inhibition of HMGB1 release(60, 61) and suppression of its cytokine and chemokine activities(62, 63).

Our current study evidenced Glycyrrhizin-mediated inhibition of acetyl-HMGB1 release to the cytoplasm in T2D/NASH conditions where acetylation of lysine residues in HMGB1 is essential for its route to the cytoplasm(64, 65). HMGB1 translocation from the nucleus to the cytoplasm in monocytes is triggered by inflammatory signals, such as lipopolysaccharide (LPS) or TNF-α, through hyper-acetylation of lysine residues within two nuclear localization sequence (NLS) sites(66). This acetylation-driven translocation is facilitated by CRM1, a key nuclear export protein(34). We have effectively showed that T2D and T2D/NASH-induced Huh7 cells have acetyl-HMGB1 translocated to the cytoplasm, but this translocation is reversed with CRM1 inhibition by LMB(67, 68). CRM1 inhibition with LMB retains CRM1 in the nucleus under T2D and T2D/NASH conditions. In hepatic cancer cells, CRM1-related transport imbalances can inactivate tumor-suppressor proteins or increase anti-apoptotic oncoproteins, aiding in cancer cell survival. This has led to the exploration of CRM1 inhibitors for cancer therapy, as blocking CRM1 function can induce cancer cell death(68, 69).HMGB1 interaction with CRM1 is significant in hepatocytes under oxidative stress a common condition given the liver role in detoxification and its frequent exposure to inflammatory stimuli(70). Modified HMGB1 binds to CRM1 for active export to the cytoplasm, where it can then move to the extracellular space. Extracellularly, HMGB1 acts as a damage-associated molecular pattern (DAMP) and binds to receptors like RAGE and TLR4 on immune cells, initiating inflammation. This inflammatory role of HMGB1 is especially relevant in liver diseases like fibrosis and hepatitis, where HMGB1 release into the extracellular space amplifies inflammation(71). Our group’s previous clinical study found elevated levels of the proinflammatory acetylated form of HMGB1 in chronic T2D patients associated with vascular complications(72).

Our study has some limitations. Our main objective aimed to assess the distribution of the nuclear transporter CRM1 in liver tissues using immunohistochemistry. We were able to detect CRM1 protein in the nuclear and cytoplasmic compartments, although the signal was weak in some samples. CRM1 immunohistochemistry-positive cells showed some sections had weaker CRM1 staining due to biological variability but had a small impact in nuclear localization in hepatocytes. Our evaluation of CRM1/HMGB1 distribution in the liver sections from human T2D or T2D/NASH subjects is a steppingstone to delineate the molecular mechanism of CRM1/HMGB1 in the preclinical T2D/NASH animal model. However, an important topic for discussion is our relatively small sample size. Even though we have a small number of samples per group that limits generalizability, our data is promising and illustrates a margin that would be generally increased with a larger cohort since we also found some differences in our histological liver damage severity among T2D and T2D/NASH patients. Such pathobiological variability shows heterogeneity due to the stratification and degree of severity from each patient.

Finally, we want to discuss our analysis of vascular vs hepatic specific source of acetyl-HMGB1. This is particularly important because it is hard to determine whether the effect evidenced in our study corresponds specifically to immune cell sources of HMGB1 being released or in either vascular or hepatic regions of interest. The observed results in isolated hepatocytes, support a hepatic contribution stronger than in immune cells in vascular rich areas. Further cell-specific analyses would better clarify these contributions in which we look to underpin the contribution of hepatocyte specific release of HMGB1 via secondary necrosis or apoptosis pathways. Such complexities arise even with our in vitro model of T2D and NASH. These models may not fully capture the complexity of human physiology since cell-based approaches have limitations in representing systemic disease processes. We are looking to advance these approaches in our future studies using in vivo models or co-culture systems that may provide more comprehensive insights.

Our findings provide further evidence of the key mechanistic role of HMGB1 silencing in the liver. We had previously shown the significant reduction in blood glucose levels improved β-cell morphology and decreased pancreatic islet inflammation underscores HMGB’’s impact on metabolic and inflammatory pathways in diabetes. In addition, decreased proinflammatory markers (TNF-α and IL-6) and decreased insulin resistance were associated with T2D HMGB1 KO mice restoring glucose.

In conclusion, our study highlights how immune cell infiltration, proinflammatory acetyl-HMGB1 translocation to cytoplasm, and CRM1 nuclear localization are key in the pathophysiology of T2D and T2D/NASH related liver damage. Unbiased examination confirmed the steatosis with increased granulocyte infiltration in T2D/NASH subjects. Acetyl-HMGB1 translocation from the nucleus to the cytoplasm was more pronounced in T2D/NASH subjects, indicating an exacerbated inflammatory state in these individuals. Furthermore, nuclear protein CRM1 showed impaired nuclear translocation in T2D/NASH liver section, possibly contributing to the inefficient export of acetyl-HMGB1. Pharmacological inhibition of CRM1 by LMB in Huh7 cells limits cytoplasmic translocation of acetyl-HMGB1 and Glycyrrhizin inhibited proinflammatory acetyl-HMGB1 might reduce liver inflammation cascade. In conclusion, these findings suggest that nuclear exporter protein CRM1 and proinflammatory protein acetyl-HMGB1 are critical players of inflammation in T2D/NASH subject liver section and Huh7 human hepatocytes and targeting their mechanism may deliver the therapeutic potential for non-alcoholic steatosis liver disease.

## Availability of Data and Materials

The data that support the findings of this is available in the MENDELEY DATA repository: Mota Alvidrez, Roberto (2024), “Targeting CRM1-HMGB1 Nuclear Translocation in Type 2 Diabetes and Metabolic Dysfunction Associated Steatotic Disease”, Mendeley Data

## Author Contributions

PP: Writing – review & editing, Writing – original draft, Software, Methodology, Investigation, Formal analysis, Data curation. BC, GMB: Validation, Resources, Methodology, Formal analysis, Data curation. JR, SPM: Visualization, Validation, Supervision, Resources, Methodology, Funding acquisition, Conceptualization. RIMA: Writing – review & editing, Writing – original draft, Visualization, Validation, Supervision, Software, Resources, Project administration, Methodology, Investigation, Funding acquisition, Formal analysis, Data curation, Conceptualization. All authors contributed to editorial changes in the manuscript. All authors read and approved the final manuscript.

## Competing Interest

The authors declare that no significant conflicts of interest exist.

## Funding

NIGMS 3R35GM119526-07S1 to RIMA, Department of Surgery and Pittsburgh Liver Research Center P&F Award Funding for RIMA (P30 DK120531), NHLBI R25HL145817 to RIMA, NCATS KL2 TR001448 funding for RIMA (UNM HSC CTSC).

## Acknowledgements

We thank Dr. Paul Monga at Pittsburgh Liver Research Center from University of Pittsburgh for sharing formalin-fixed paraffin-embedded human liver biopsies. We would like to thank Dr. Sharina Desai and Dr. Li Chen for their invaluable assistance with technical issues related to the use of the CX7 Cellomics imaging platform. We thank Cathy Martinez and Fred Schultz from the Human Tissue Repository for their support with liver histology and immunohistochemistry. We would like to express our sincere gratitude to Dr. Jaya Rajaiya and her team providing invaluable resources, expertise, and support throughout this study.

---

**Supplementary Figure 1. Human liver tissue histological analysis using hematoxylin and eosin staining from control, T2D and T2D/NASH liver tissues focused in vascular areas.** Control group: normal histological morphology and no steatosis; T2D group shows infiltration of blood cells and steatosis grade 2 and stage 1; T2D/NASH group; histological section appears stage 4 and grade 1 steatosis (A). Agranulocytes and granulocytes were analyzed and expressed as follows: total cell per tissue area (B), average cytoplasm per cell area (C), and average nucleus per total cell area(D). All data represented mean+/-SEM. All granulocytes and Agranulocytes numbers in control compared to T2D or T2D/NASH of humans (n=4) are used for One-way ANOVA analysis. Significance levels: *:<0.05, **: 0.001-0.01, ***: 0.0001, ****: <0.0001.

**Supplementary Figure 2. Hematoxylin as a nuclear counter stain to distinguish between cytoplasmic and nucleus of hepatic and vascular region of masked and unmasked region of acetyl-HMGB1 and CRM1 using HALO software.** Hepatic region: unmasked acetyl-HMGB1(A), and masked acetyl-HMGB1 (B). and Vascular region: unmasked acetyl-HMGB1(C), masked acetyl-HMGB1 (D); Hepatic region: unmasked CRM1(E), masked CRM1(F), and Vascular region: unmasked CRM1(G), masked CRM1 (H).

**Supplementary Figure 3. Hepatic acetyl-HMGB1 protein in nucleus and cytoplasm of liver tissue from control, T2D, and T2D/NASH human patients subjects with special ROI in vascular areas.** Formalin-fixed paraffin-embedded human liver biopsies from control, T2D, and T2D/NASH human patient samples were used for the analysis of acetyl-HMGB1 protein, which is one of the proinflammatory forms of HMGB1 to distinguish disease severity. Weak positive, moderate positive, and strong positive of acetyl-HMGB1 protein analyzed as follows: A) Nucleus, B) Cytoplasmic, C) quantification of cytoplasmic/nuclear, D) quantification of nuclear/cytoplasmic acetyl HMGB1. All data represented mean+/-SEM. All nuclear and cytoplasmic acetyl-HMGB1 protein in control compared to T2D or T2D/NASH of humans (n=4) are used for One-way ANOVA analysis. Significance levels: *:<0.05, **: 0.001-0.01, ***: 0.0001, ****: <0.0001. Immunohistochemistry of acetyl-HMGB1 protein color shows as brown and captured at x400.

**Supplementary Figure 4. Hepatic CRM1 protein in nucleus and cytoplasm of control, T2D, and T2D/NASH human patients subjects with special ROI in vascular areas.** Formalin-fixed paraffin-embedded human liver biopsies from control, T2D, and T2D/NASH human patient samples were used for the analysis of CRM1 protein, which is one of the nuclear exporter proteins. Weak positive, moderate positive, and strong positive of CRM1 protein expression analyzed as follows: A) Nuclear, B) cytoplasmic, C) quantification of cytoplasmic/nuclear, D) quantification of nuclear/cytoplasmic of acetyl-HMGB1 protein expression. All data represented mean+/-SEM. All nuclear and cytoplasmic acetyl-HMGB1 in control compared to T2D or T2D/NASH of humans (n=4) are used for One-way ANOVA analysis. Significance levels: *:<0.05, **: 0.001-0.01, ***: 0.0001, ****: <0.0001. Immunohistochemistry of CRM1 protein color shows as brown and captured at x400.

**Supplementary Figure 5. Non-T2D (control) Huh7 hepatocyte HMGB1 in the cytoplasm and nucleus treated with CRM1 inhibitor (LMB).** In the control cells treated with or without 25nM of Leptomycin B for 12hrs then nuclear and cytoplasmic proteins were isolated for western blotting. Cytoplasmic and nuclear acetyl-HMGB1 and CRM1 proteins were analyzed using western blot (A), Cytoplasmic acetyl-HMGB1(B) and CRM1(C), and nuclear acetyl-HMGB1(D) and CRM1(E). All data represented mean+/-SEM. All nuclear and cytoplasmic acetyl-HMGB1 and CRM1 protein in non-T2D compared to Non-T2D with LMB (n=3) are used for student t-test analysis. Significance levels: *:<0.05, **: 0.001-0.01, ***: 0.0001, ****: <0.0001.

**Supplementary Figure 6. Non-T2D (control) Huh7 hepatocyte HMGB1 in the cytoplasm and nucleus treated with HMGB1 inhibitor (GA).** In the control cells treated with or without 50µM of Glycyrrhizin for 24hrs then nuclear and cytoplasmic proteins were isolated for western blotting. Cytoplasmic and nuclear acetyl-HMGB1 and CRM1 proteins were analyzed using western blot (A), Cytoplasmic acetyl-HMGB1(B) and CRM1(C), and nuclear acetyl-HMGB1(D) and CRM1(E). All data represented mean+/-SEM. All nucleus and cytoplasmic acetyl-HMGB1 and CRM1 protein in non-T2D compared to Non-T2D with GA (n=3) are used for student t-test analysis. Significance levels: *:<0.05, **: 0.001-0.01, ***: 0.0001, ****: <0.0001.

## Notes

### Competing Interest Statement

The authors have declared no competing interest.

